# Involvement of Brn3a-positive spinal dorsal horn neurons in the transmission of visceral pain in inflammatory bowel disease model mice

**DOI:** 10.1101/2021.09.06.457875

**Authors:** Kazuhiko Nishida, Shinji Matsumura, Takuya Kobayashi

## Abstract

The spinal dorsal horn (SDH) plays a crucial role in the transmission and processing of somatosensory information. Although spinal neural circuits that process several distinct types of somatic sensations have been studied extensively, those responsible for visceral pain transmission remain poorly understood. In the present study, we analyzed dextran sodium sulfate (DSS)-induced inflammatory bowel disease (IBD) mouse models to characterize the spinal dorsal horn neurons involved in visceral pain transmission. Immunostaining for c-fos, a marker of neuronal activity, demonstrated that numerous c-fos-positive cells were found bilaterally in the lumbosacral spinal dorsal horn, and their distribution was particularly abundant in the shallow dorsal horn. Characterization of these neurons by several molecular markers revealed that the percentage of the Pit1-Oct1-Unc86 domain (POU domain)-containing transcription factor Brn3a-positive neurons among the c-fos-positive neurons in the shallow dorsal horn was 30-40% in DSS-treated mice, which was significantly higher than that in the somatic pain model mice. We further demonstrated by neuronal tracing that, within the shallow dorsal horn, Brn3a-positive neurons were more highly represented in spino-solitary projection neurons than in spino-parabrachial projection neurons. These results raise the possibility that Brn3a-positive spinal dorsal horn neurons make a large contribution to visceral pain transmission, part of which is mediated through the spino-solitary pathway.

## INTRODUCTION

Sensory signals from internal organs are crucial for monitoring visceral disorders such as inflammation or mechanical distention. Such visceral information is conveyed to higher brain centers via the spinal cord and vagus nerve, with the spinal cord pathway commonly assumed to transmit pain (1). Within the spinal cord, the dorsal horn plays a pivotal role in processing sensory signals and their transmission to the higher brain center, which are mediated by heterogeneous populations of interneurons and supraspinal projection neurons, respectively. Several studies on spinal sensory transmission indicate that distinct neuronal populations process different somatosensory modalities, such as mechanical pain, heat, and itch (2). Visceral pain is recognized as a sensation that differs from somatic pain (e.g., diffused pain localization and occasional induction of referred pain and autonomic reflex) (3). Visceral and somatic pain may be processed by distinct neuronal circuits in the spinal dorsal horn. This notion has been confirmed by previous studies on neuronal activity patterns in the spinal dorsal horn in response to visceral and somatic stimulation. Visceral stimulation evokes the activity of spinal neurons primarily in the marginal (lamina I) and deeper (lamina V and X) laminae, as well as in the parasympathetic nucleus (4-7). In contrast, somatic stimulation induces spinal neurons broadly distributed across the laminae of the spinal dorsal horn (lamina I-V) (4, 5). There have been controversial reports regarding the response specificity of the spinal dorsal horn neurons. Cervero et al. reported that most spinal dorsal horn neurons in lamina I and V responsive to visceral stimulation also received somatic sensory inputs (4, 8). In contrast, Luz et al. recently showed that lamina I neurons in the spinal dorsal horn comprise visceral-, somatic-, and viscero-somatic-responsive neurons (9), suggesting that somatic and visceral pain might be processed by distinct neuronal circuits. The ascending spinal pathway, necessary for visceral pain transmission to the higher brain center, has been the subject of several studies. A population of supraspinal projection neurons in the marginal lamina responds to visceral stimulation but not to somatic stimulation (9). In addition, postsynaptic dorsal column (PSDC) neurons, other subtypes of supraspinal projection neurons, have been shown to play a critical role in visceral pain transmission to the gracile nucleus in the medulla (10, 11). Although we know more about the distribution pattern and morphological features of visceral pain-responsive neurons in the spinal dorsal horn, we still know very little about the neuronal identity of these neurons, which hampers the understanding of spinal neural circuits for visceral pain transmission.

Brn3a, a Pit1-Oct1-Unc86 domain (POU domain)-containing transcription factor, is expressed in a wide variety of long-range neurons in the central and peripheral nervous systems, including dorsal root ganglion cells, retinal ganglion cells, and medial habenula neurons (12). The expression of Brn3a in the developing spinal cord has long been known (13, 14), and Brn3a-positive neurons are eventually localized in the marginal and deeper dorsal horns at postnatal stages (15). However, their functions in sensory processing remain unclear.

In the present study, we employed dextran sodium sulfate (DSS)-treated inflammatory bowel disease (IBD) models as a possible model for visceral pain. Analysis of neuronal activity in the lumbosacral spinal cord revealed that Brn3a-positive neurons in the marginal lamina were preferentially activated under visceral pain conditions compared with somatic neurons. We also showed that Brn3a-positive neurons responsive to visceral pain are involved in viscerosensory transmission to the nucleus of the solitary tract (NTS).

## METHODS

### Animals

C57BL/6J mice (CLEA Japan Inc., Tokyo, Japan) were maintained on a 12 h (8:00 to 20:00) light-dark schedule. All animal experiments were conducted in accordance with the Japanese guidelines and regulations (2006) for scientific and ethical animal experimentation and were approved by the Animal Experimentation Committee of Kansai Medical University (21-037).

### Induction of Experimental Colitis

Seven- to nine-week-old male mice were treated with drinking water containing 2% DSS (molecular weight: 36,000 - 50,000, MP Biomedicals, Illkirch, France) or regular drinking water as a control. Mice were treated with DSS-containing water for 7-8 days until their body weight was reduced by 10-20% compared to that before treatment. The behavioral test was performed six days after DSS treatment. For the behavioral test, DSS-treated and control mice were placed on a wire mesh enclosed by an acrylic wall (10 cm in length and width and 25 cm in height). Following habituation for 20 min, the behavior of the mice was videotaped for 30 min. The duration and number of licks on the anus were analyzed.

### Histological Analysis of the Colon

The DSS-treated and control mice were transcardially perfused with 0.1 M phosphate buffer (pH 7.4) containing 4% paraformaldehyde following anesthesia with pentobarbital (50 mg/kg, Nacalai Tesque, Kyoto, Japan). The colons were dissected, slit opened with scissors, and rolled up using a wooden stick, as described previously (16). Following post-fixation overnight, the samples were immersed overnight at 4°C in 0.1 M phosphate-buffered saline (PBS) containing 30% (w/w) sucrose. The samples were then embedded in the OCT compound (Sakura Finetek, Tokyo, Japan) and stored at −80°C until further examination. The 10-μm sections of the colon were stained with hematoxylin and eosin (HE), and the images were then obtained using a Nano-Zoomer slide scanner (Hamamatsu Photonics K.K., Hamamatsu, Japan).

### Induction of Cutaneous Pain by Formalin Injection

To apply cutaneous pain to the skin, formalin was injected into the lower back skin on the right side, which is between the roots of the tail and the lower limb of the mouse. Primary sensory neurons responsible for this skin area project to the spinal dorsal horn at the L6 – S1 segments (17), where visceral pain inputs in DSS-treated mice were found in the present study. Hair from the skin around this area was removed using an electric shaver one day before injection. Following brief anesthesia with isoflurane, 10 μl of 5% formalin was injected into the skin using a 30-gauge needle attached to Hamilton syringe. Two hours after injection, the mice were perfused to obtain spinal cord samples.

### Immunohistochemistry and *in situ* Hybridization

The DSS-treated and formalin-injected mice were transcardially perfused with 0.1 M phosphate buffer (pH 7.4) containing 4% paraformaldehyde following anesthesia with pentobarbital (50 mg/kg), and their spinal cords were subsequently dissected out. Following post-fixation for 3 h, the samples were immersed overnight at 4°C in 0.1 M PBS containing 30% (w/w) sucrose. Before the sample was frozen, spinal cord samples were notched so that the L4 – S1 spinal segments could be distinguished in later analyses. The spinal cords were then embedded in the OCT compound and stored at −80°C until further examination.

Transverse sections of the lumbosacral spinal cord of the mice were prepared using a cryostat (20 μm sections). Immunohistochemistry of the spinal dorsal horn sections was performed as previously `described` (18). Antibodies used in the current study are goat anti-c-fos (1:200, Santa Cruz Biotechnology, Dallas, TX, USA, sc-52), rabbit anti-c-fos (1:500, Synaptic Systems, Goettingen, Germany, 226003), mouse anti-Brn3a (1:300, Sigma, St. Louis, MO, USA, MAB1585), rabbit anti-Pax2 (1:500, Thermo Fisher Scientific Inc., Waltham, MA, USA, 71-6000), goat anti-cholera toxin B (CTB, 1:5,000, List Biological Laboratories, Campbell, CA, USA, 703), rabbit anti-PKCγ (1:100, Santa Cruz Biotechnology, sc-211), Alexa 594-conjugated donkey anti-goat IgG (1:300, Invitrogen, Carlsbad, CA, USA, A11058), Alexa 594-conjugated goat anti-mouse IgG (1:300, Invitrogen, A11032), Alexa 488-conjugated donkey anti-mouse IgG (1:300, Invitrogen, A21202), Alexa 488-conjugated donkey anti-rabbit IgG (1:300, Invitrogen, A21206), and Cy5-conjugated donkey anti-mouse IgG antibodies (1:300, Jackson ImmunoResearch, West Grove, PA, USA, 715-175-150).

*In situ* hybridization coupled with immunostaining was performed as previously described (19), with some modifications. cDNAs for mouse *somatostatin* (*SST*, 15-528 of NM_009215), mouse *preprodynorphin* (*Pdyn*, 1-747 of NM_018863), and mouse *cholecystokinin* (*CCK*, 3-688 of NM_031161) were PCR-amplified and subcloned into the pGEM-T Easy vector (Promega, Fitchburg, WI, USA). Digoxigenin (DIG)-labeled anti-sense probes were prepared using DIG RNA Labeling Mix (Sigma, 11277073910), SP6 RNA polymerase (Sigma, 10810274001), and template cDNAs and purified using SigmaSpin Sequencing Reaction Clean-Up (Sigma, S5059). Transverse sections of the spinal cord were post-fixed for 15 min, washed with diethylpyrocarbonate (DEPC)-treated dH_2_O, and incubated with 0.3% H_2_O_2_ for 10 min to inactivate endogenous peroxidases. After washing with DEPC-treated dH_2_O, they were incubated for 10 min with 0.1 M triethanol amine (pH 8.0) containing 0.25% acetic anhydride. The sections were then incubated at 60°C for 1 h in a hybridization buffer containing 50% formamide, 5 × SSPE (pH 7.4), 1 × SDS, and 0.4 mg/ml yeast tRNA (Sigma, 10109517001). They were then transferred to a hybridization buffer containing a DIG-labeled riboprobe that had been heat-denatured at 80°C for 5 min in advance and kept at 60°C overnight. The sections were washed three times at 60°C for 30 min with hybridization wash buffer (50% formamide, 2 × SSC [pH 7.0]) and subsequently three times at room temperature for 5 min with Tris-buffered saline (TBS, pH 7.5) containing 0.1% Tween-20. After blocking with TSA blocking reagent (PerkinElmer, Waltham, MA, USA, FP1012) for 1 h, sections were incubated with TSA blocking reagent containing peroxidase-conjugated sheep anti-DIG antibody (1:1,000, Sigma, 11207733910) at room temperature for 2 h. After washing with TBS containing 0.1% Tween-20, the sections were incubated with Tyramide-conjugated Cy3 or fluorescein (1:75, PerkinElmer, NEL744001KT, NEL741001KT) at room temperature for 20 min to visualize the *in situ* signals. Then, after washing with PBS containing 0.2% Triton X-100, they were further incubated with a blocking solution (PBS containing 0.2% Triton X-100 and 2% BSA) at room temperature for 1 h, followed by incubation with rabbit anti-c-fos antibody (Synaptic Systems) or mouse anti-Brn3a antibody (Sigma) in blocking solution at 4°C overnight. Thereafter, they were incubated with Alexa 488-conjugated donkey anti-rabbit IgG (1:300, Invitrogen) or Alexa488-conjugated donkey anti-mouse IgG (1:300, Invitrogen) to visualize c-Fos or Brn3a signals, respectively.

The sections were counterstained with Hoechst 33342 (1:2,000, Thermo Fisher Scientific Inc., H3570) and were mounted with a mounting reagent containing 25% glycerol, 0.1 M Tris (pH 8.5), 27.5 mg/ml 1,4-Diazabicyclo[2.2.2]octane (DABCO, Tokyo Chemical Industry Co., Tokyo, Japan, D0134), and 100 mg/ml Mowiol 4-88 (Merck, Darmstadt, Germany, 475904), and fluorescent images were then obtained with a confocal microscope (LSM700, Carl Zeiss, Jena, Germany).

### Retrograde Labeling of Primary Sensory Neurons and Supraspinal Projection Neurons by CTB

For retrograde labeling of primary sensory neurons responsible for the anal skin, wild-type mice were deeply anesthetized with a mixture of medetomidine (37.5 μg/kg, Nippon Zenyaku Kogyo, Fukushima, Japan), midazolam (2 mg/kg, Sandoz, Tokyo, Japan), and butorphanol (0.25 mg/kg, Meiji Seika Pharma, Tokyo, Japan), and the hair around the anus was removed using an electric shaver. 1% CTB (Sigma, C9903) was loaded into a fine glass capillary and was injected subcutaneously into three different parts of the skin around the anus on the left by a microinjector (IM-31, Narishige, Tokyo, Japan). One microliter of CTB solution was used for each injection. Four days later, the mice were treated with water containing 2% DSS for seven days to induce colitis. The mice were transcardially perfused to obtain lumbosacral spinal cord samples. Transverse sections (20 μm in thickness) of the samples were immunostained with antibodies against c-Fos and CTB.

The CTB injection into the lateral parabrachial nucleus has been previously `described` (18). A fine glass capillary (tip diameter: ∼50 μm) connected to a Hamilton syringe was used for CTB injection into the NTS. Following anesthesia with a mixture of medetomidine, midazolam, and butorphanol, 7- to 9-week-old mice were fixed in a stereotactic apparatus (SR-5M-HT, Narishige, Tokyo, Japan) and a burr hole was made in the skull of the mice over the injection site (bregma −8.0, lateral 0) using a dental drill. A glass capillary filled with 1% CTB (Sigma) was inserted into the brain 4.0 mm below the brain surface, and 400 nl of CTB solution was injected into the brain using a microinjector (IMS-20, Narishige) attached to a stereotactic apparatus. Six days after the CTB injection, the mice were transcardially perfused to obtain spinal cord and hindbrain samples. Since the NTS is localized around the midline of the medulla, CTB injection labeled both the right and left sides of the nucleus. Thus, retrogradely labeled lamina I spino-solitary neurons on both sides were analyzed in this study.

For labeling of spino-solitary supraspinal projection neurons in DSS-treated mice, CTB was first injected into NTS, and treatment with 2% DSS started 2 days later. Nine to ten days after the CTB injection, the mice were transcardially perfused to obtain spinal cord and hindbrain samples for further analysis.

### Image Analysis

To analyze the global distribution pattern of c-fos-positive cells in control, DSS-treated, and formalin-injected mice, several fluorescence images covering the entire spinal cord section were taken using a confocal microscope (LSM700) and stitched together to obtain whole images. Darkfield photographs were also taken to distinguish the boundary between the white and gray matter. ImageJ was used to measure the fluorescence intensity. Transverse sections of the spinal cord were divided into four regions in this study: shallow dorsal horn (SDH), deep dorsal horn (DDH), ventral horn (VH), and lateral funiculus (LF). Dorsal and ventral horns were defined as areas of gray matter dorsal and ventral to the central canal, respectively. The dorsal horn was further divided into the SDH and DDH by a line between the ventral edge of the dorsal funiculus and the medial edge of the LF in the intermediate spinal cord.

For the analysis of the signal intensity of several markers among c-fos-positive or Brn3a-positive neurons, fluorescence images were captured every 2 μm along the z-axis, and the average fluorescence intensity was measured using Imaris (Oxford Instruments, Abington-on-Thames, UK). The c-fos-positive neurons with average c-fos signal intensity above 5,000 and 10,000 were defined as c-fos and c-fos-H cells, respectively. For the quantification of Pax2 and Brn3a signals, the average intensity, which was two times stronger than the background level, was regarded as a positive signal. For the quantification of somatostatin and preprodynorphin signals, the average intensity, which was three times stronger than the background level, was regarded as a positive signal.

### Statistical Analyses

The results of animal behavior and histochemistry were analyzed using the Mann-Whitney U test. All statistical analyses were performed using GraphPad Prism 7.02 (GraphPad Software Inc., La Jolla, CA, USA). Data are expressed as mean ± SEM.

## RESULTS

### Generation of DSS-Induced Colitis as a Visceral Pain Model

Inflammation of the gastrointestinal tract is caused by IBD, which is often accompanied by chronic abdominal pain in humans (20). To investigate the spinal neuronal circuits involved in visceral pain transmission, we induced IBD in mice by treatment with DSS-containing water as a possible model for visceral pain. C57BL/6J mice were used in this study because their sensitivity to DSS treatment is sufficiently high to cause severe inflammation (21) and evoke neuronal activity in the SDH (22). Mice treated with 2% DSS exhibited occult blood in their stool and diarrhea from around the 4^th^ day, followed by body weight loss (Figure. 1A). Treatment with DSS for seven days induced mucosal damage, mainly in the distal rather than the proximal part of the colon (Supplementary Figure 1), as reported previously (23). Licking the body surface close to the visceral organ is a common spontaneous behavior in several visceral pain models (24-26). Therefore, we assessed the licking behavior of mice treated with DSS using video recordings. This analysis demonstrated that these mice exhibited increased number (Pre: 1.1 ± 0.7 [*n* = 7 mice], Post: 12.6 ± 2.7 [*n* = 7 mice]) and duration (Pre: 7.3 ± 4.4 s [*n* = 7 mice], Post: 100.1 ± 17.4 s [*n* = 7 mice]) of licking around the anus post-DSS-treatment compared with pre-DSS-treatment (Figure 1B and 1C, Supplementary Figure 2). Daily observations of this behavior following DSS treatment showed that the onset of licking often precedes that of diarrhea (Supplementary Figure 2, left), suggesting that this behavior is not dependent on diarrhea, but rather on inflammation-evoked visceral pain. These results suggest that DSS treatment is a valid method for generating a mouse model of visceral pain.

**FIGURE 1.**
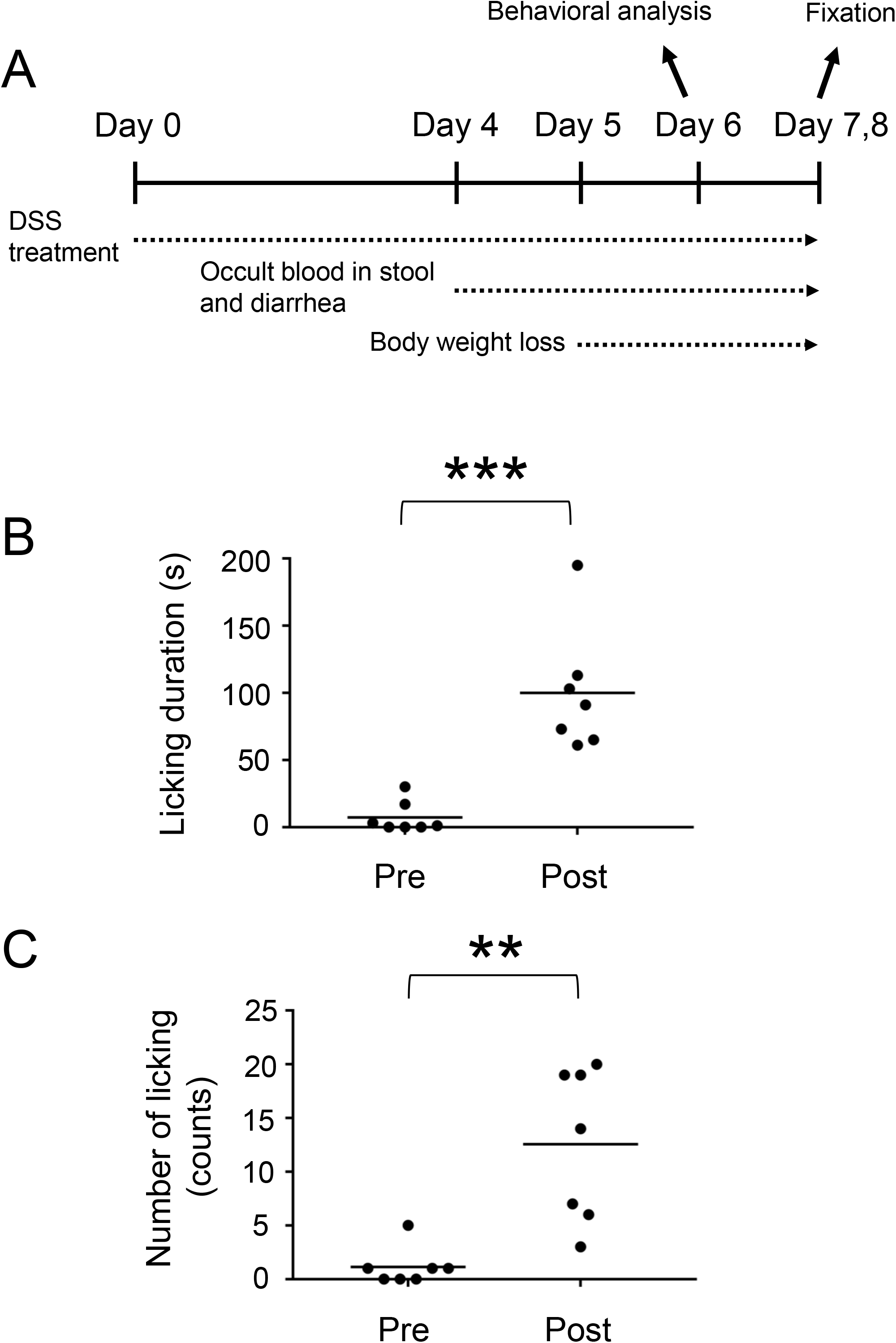
DSS-treated mice as a visceral pain model. **(A)** Experimental design of DSS-treated mice. Duration **(B)** and number **(C)** of licking behavior during a 30 min time window before (Pre) and after (Post) treatment with 2% DSS are shown. Horizontal bars indicate the median. Data were analyzed by the Mann-Whitney U test. **p < 0.01, ***p<0.001.

### Neural Activity in the Spinal Cord of DSS-Treated Mice

To directly assess whether the spinal dorsal horn receives visceral sensory inputs in mice treated with DSS, we analyzed the global distribution of neuronal activity in the spinal cord of mice by c-fos immunostaining. For this analysis, DSS-treated mice with a 10 to 20 % decrease in body weight compared to pretreatment time were used. The reduction in body weight to this extent typically took 7–8 days after DSS treatment (Figure 1A). Mice in this condition are hereafter referred to as “DSS-treated mice”. Noxious sensation in the distal colon is transmitted via lumbar splanchnic and pelvic afferents, whose central terminals are localized at the thoracolumbar (T10-L1) and lumbosacral (L6-S1) spinal cords, respectively (27). The c-fos immunostaining of the spinal cord sections of DSS-treated mice demonstrated that numerous c-fos-positive neurons were found in the lumbosacral spinal cord compared with the control mice, with the highest abundance between the L6 and S1 spinal segments (Figure 2). However, we could hardly find c-fos-positive neurons in the rostral spinal segments, including T10 to L1 (Supplementary Figure 3). These results suggest that visceral pain in DSS-treated mice is mainly transmitted through the pelvis but not the lumbar splanchnic nerves.

**FIGURE 2.**
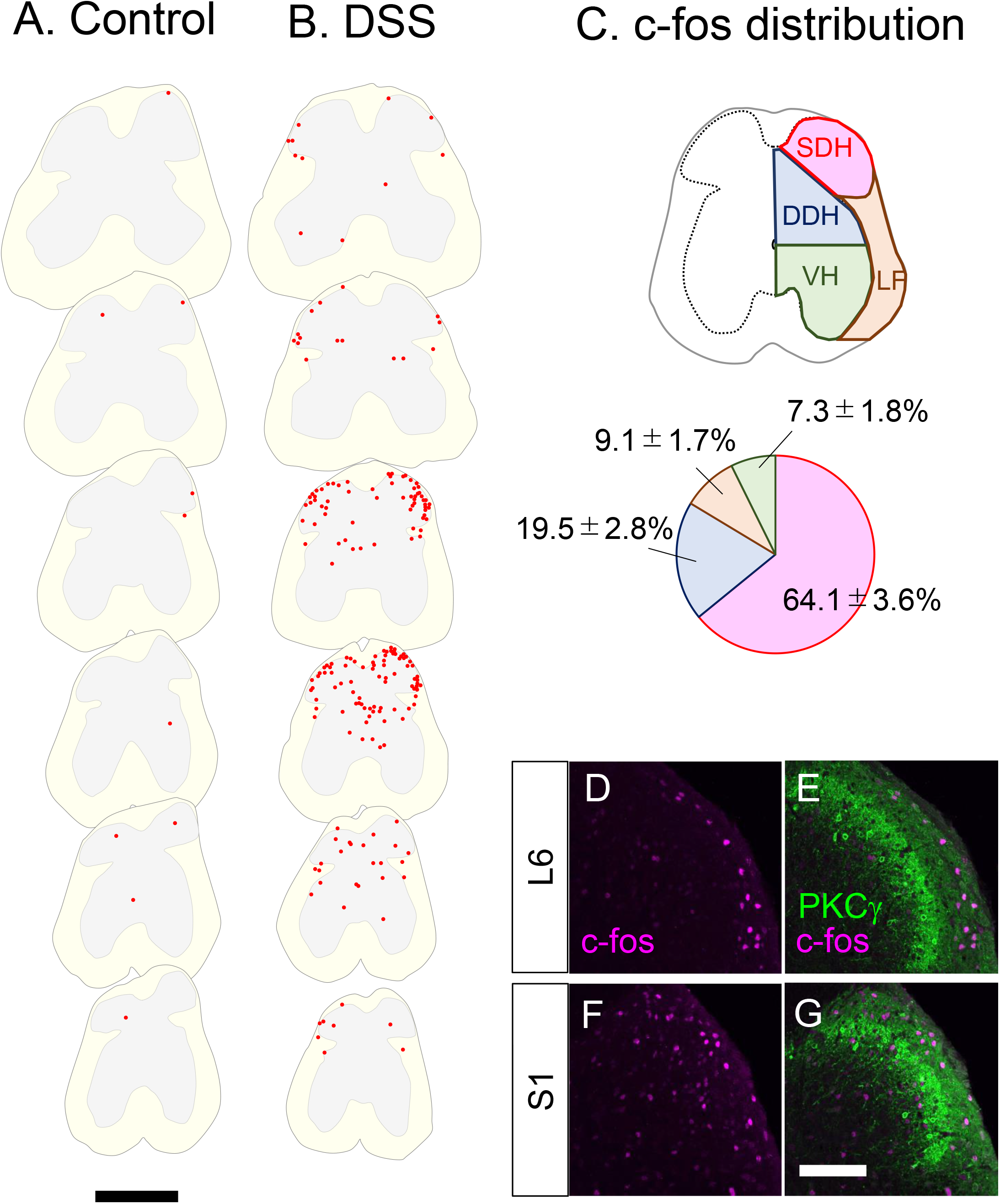
c-fos-positive cells in the lumbosacral spinal cord in control and DSS-treated, mice. **(A, B)** Distribution of c-fos-positive cells. c-fos immunostaining on the transverse sections of the lumbosacral spinal cords in 0% DSS-treated **(A)** and 2% DSS-treated **(B)** mice was performed, and distribution of c-fos-positive cells on six sections (20 μm in thickness, 800 μm apart each other) from the lumbar (∼L5, top) to the sacral (∼S2, bottom) spinal cords in a representative mouse in each condition is shown. c-fos-positive cells are shown as red dots. White and gray matters in each section are shown in light yellow and grey colors, respectively. Scale, 500 μm. **(C)** Distribution pattern of c-fos-positive cells. The spinal cord was divided into the shallow dorsal horn (SDH), deep dorsal horn (DDH), ventral horn (VH), and lateral funiculus (LF) (see Materials and Methods). The distribution of c-fos-positive cells in these areas in DSS-treated mice (*n* = 4 mice, 1,205 cells) is shown in the pie chart. **(D-G)** Transverse sections of the spinal dorsal horn of DSS-treated mice were immunostained with anti-c-fos (magenta) together with anti-PKCγ (green) antibodies. Images on the right dorsal horn around L6 **(D, E)** and S1 **(F, G)** segments are shown. Scale, 100 μm.

Further analysis of c-fos distribution in the lumbosacral spinal cord of DSS-treated mice demonstrated that 64.1 ± 3.6% (*n* = 4 mice, 1,205 cells) of c-fos-positive neurons in DSS-treated mice were found in the SDH, with the remainder distributed in the DDH, LF, and VH (Figure 2C). The abundance of c-fos-positive neurons in the SDH prompted us to further analyze their detailed distribution within the SDH. To this end, we analyzed the laminar localization of c-fos-positive neurons in the SDH by co-immunostaining with PKCγ (Figure 2D-G), a marker of neurons at the lamina II-III border (2). Within the SDH, c-fos-positive neurons were commonly found in the marginal lamina (lamina I) throughout segments, with some of those scattered in lamina IIo to III in the more caudal segments (∼S1). Taken together, these results suggest that visceral pain signals in DSS-treated mice are transmitted mainly to the marginal SDH of the spinal cord around the L6 to S1 segments.

### Characterization of c-fos-Positive Neurons in DSS-Treated Mice

Characterization of visceral pain-responsive neurons in the spinal cord would provide clues for understanding the neural circuits for visceral pain transmission. Thus, we characterized c-fos-positive neurons in DSS-treated mice by co-immunostaining for several markers (Figure 3A-I). Somatostatin (SST) marks ∼60% of excitatory interneurons in laminae I-II of the spinal dorsal horn (28). We performed immunostaining of c-fos together with *in situ* hybridizations of *SST* (Figure 3A-C) and demonstrated that 17.6 ± 1.7% (*n* = 6 mice, 573 cells) of c-fos-positive cells were *SST*-positive. Among those with strong c-fos intensity (c-fos-H), 21.2 ± 2.2% (*n* = 6 mice, 230 cells) were *SST*-positive. Next, we performed immunostaining for Pax2 (Figure 3D-F), a marker of GABAergic inhibitory interneurons throughout the spinal dorsal horn (29). This analysis demonstrated that 31.8 ± 1.4% (*n* = 5 mice, 1,548 cells) of c-fos-positive cells and 42.0 ± 1.4% (*n* = 5 mice, 488 cells) of c-fos-H-positive mice were Pax2-positive, which were comparable to the abundance of inhibitory neurons in the SDH (30). The preferential distribution of c-fos-positive neurons in the marginal lamina of the spinal dorsal horn in DSS-treated mice (Figure 2D-G) prompted us to characterize these neuronal populations using a marker of marginal lamina neurons. Brn3a is expressed in the marginal lamina and deeper laminae of the spinal dorsal horn at both developmental and adult stages (15), and can be used as a marker of marginal lamina neurons in the SDH. Thus, we performed immunostaining using antibodies against c-fos and Brn3a in DSS-treated mice (Figure 3G-I). This analysis showed that 30.4 ± 2.6% (*n* = 5 mice, 1,141 cells) of c-fos-positive cells and 38.1 ± 5.3% (*n* = 5 mice, 324 cells) of c-fos-H-positive ones were Brn3a-positive.

**FIGURE 3.**
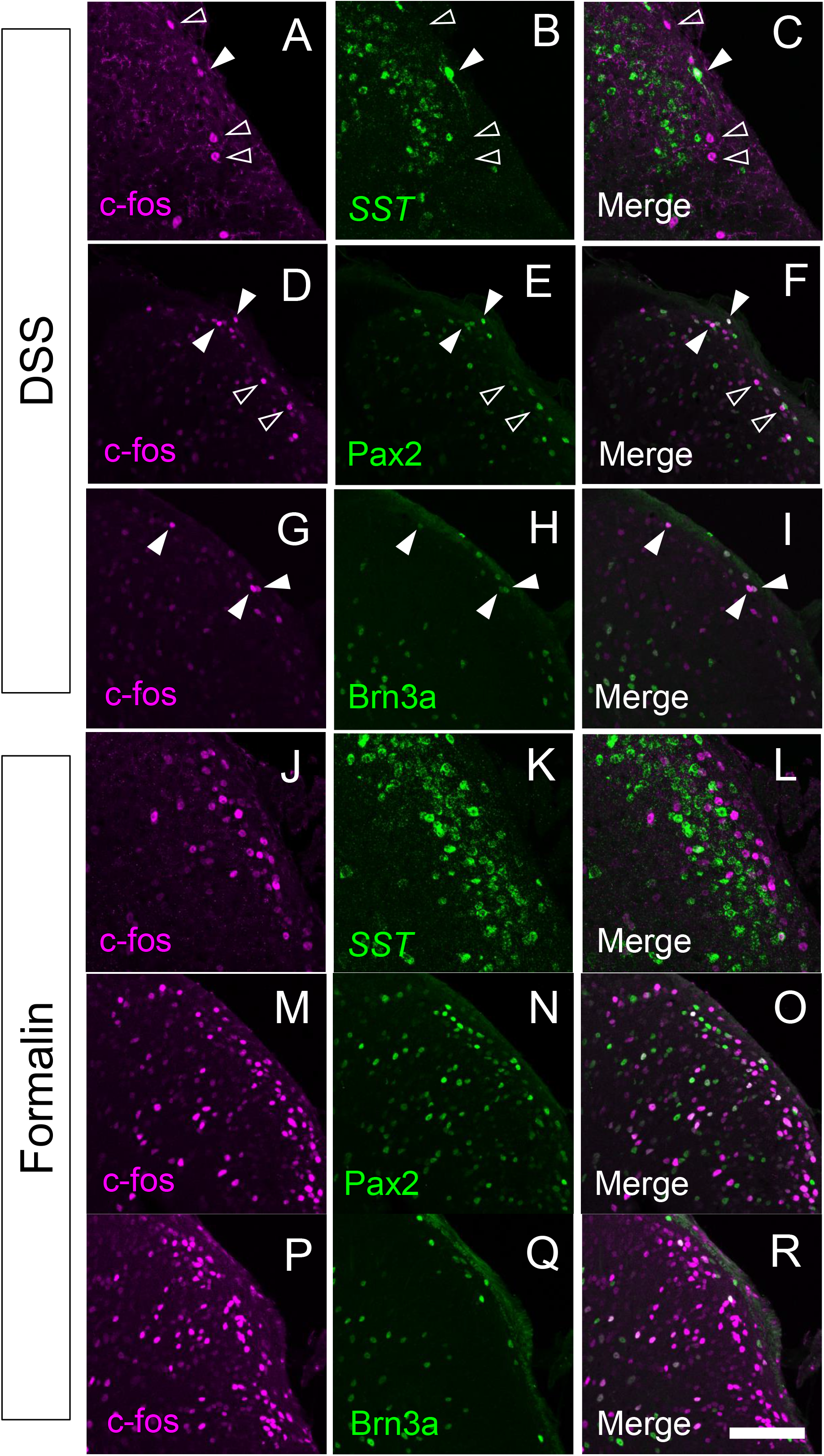
Immunostaining of c-fos together with several molecular markers in the spinal cord of DSS-treated and formalin-injected mice. c-fos immunostaining together with *somatostatin* (*SST*) *in situ* hybridization **(A-C, J-L)**, or Pax2 **(D-F, M-O)** or Brn3a **(G-I, P-R)** immunostaining were performed on transverse sections of the lumbosacral spinal dorsal horn of DSS-treated **(A-I)** and formalin-injected **(J-R)** mice. Images of c-fos immunostaining (magenta; **A, C, D, F, G, I, J, L, M, O, P, R**), *SST in situ* hybridization (green; **B, C, K, L**), Pax2 immunostaining (green; **E, F, N, O**), or Brn3a immunostaining (green; **H, I, Q, R**) on the right spinal dorsal horn are shown. Open arrowheads indicate c-fos single-positive cells, whereas closed arrowheads indicate cells positive for both c-fos and marker molecules. Scale, 100 μm.

To understand the differences in neural circuits mediating visceral versus somatic pain, we compared our visceral pain model with somatic pain model mice. The somatic pain model was generated by subcutaneous injection of formalin into the lower back skin so that somatic pain signals were introduced into the L6 to S1 spinal segments (see Materials and Methods for details). The overall distribution of c-fos positive neurons in formalin-injected mice was comparable to that in DSS-treated mice, except that c-fos-positive neurons were distributed almost exclusively on the formalin-injected side and that many of them were distributed in both laminae I and II in the SDH throughout the spinal segments (Supplementary Figure 4). Thus, we examined the expression of SST, Pax2, and Brn3a in c-fos-positive neurons of formalin-injected mice (Figure 3J-R). The percentage of *SST-* and Pax2-positive neurons among c-fos-positive neurons in formalin-injected mice was comparable to that in DSS-treated mice (Figure 4A, left and 4B, left). In contrast, the percentage of Brn3a-positive neurons among c-fos-positive neurons was significantly higher in DSS-treated mice than in the formalin model (Figure 4C, left). Similar results were obtained for the c-fos-H-positive populations (Figure 4A-C, right). These results raised the possibility that Brn3a-positive neurons in the marginal lamina make a larger contribution to the transmission of visceral pain in DSS-treated mice than somatic pain in formalin-injected mice.

**FIGURE 4.**
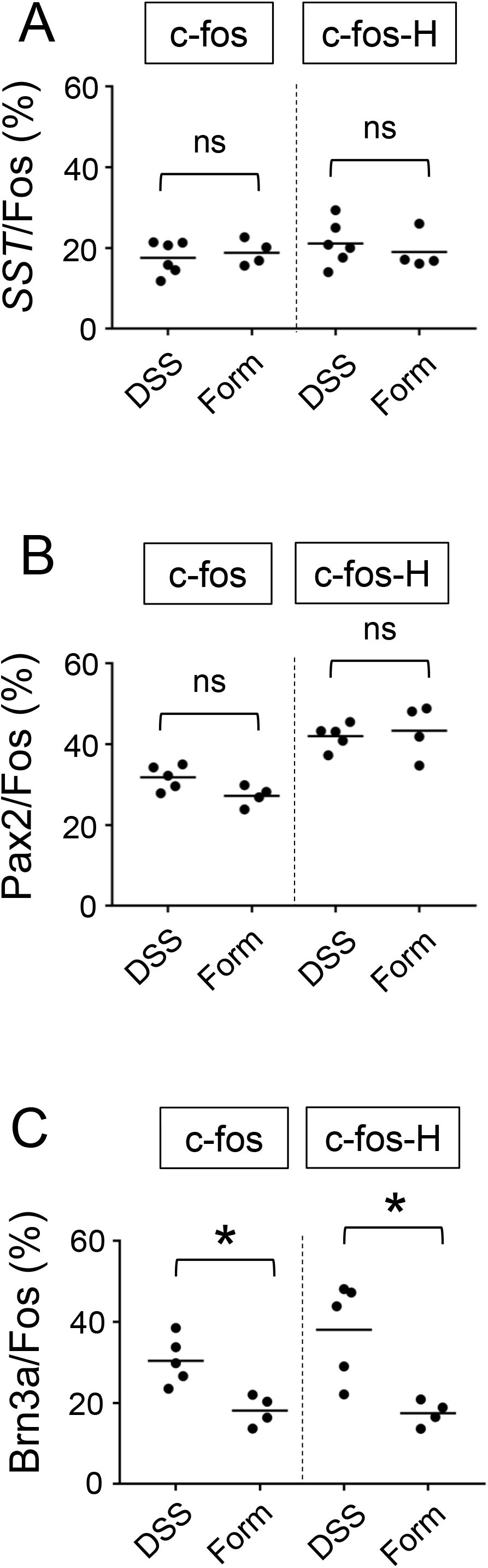
Percentage of *SST*-, Pax2-, and Brn3a-positive cells among c-fos-positive cells. c-fos-positive cells were classified into two categories: “c-fos-H” cells include those with strong c-fos intensity, whereas c-fos cells include those with both weak and strong c-fos intensity (see Materials and Methods). **(A)** The percentage of *SST*-positive cells among c-fos cells (17.6 ± 1.7%, *n* = *6* DSS-treated mice, 573 cells; 18.9 ± 1.6%, *n* = 4 formalin-injected mice, 758 cells) and c-fos-H cells (21.2 ± 2.2%, *n* = 6 DSS-treated mice, 230 cells; 19.0 ± 2.3 %, *n* = 4 formalin-injected mice, 421 cells) is shown. **(B)** The percentage of Pax2-positive cells among c-fos cells (31.8 ± 1.4% *n* = 5 DSS-treated mice, 1,548 cells; 27.2 ± 1.3%, *n* = 4 formalin-injected mice, 1,484 cells) and c-fos-H cells (42.0 ± 1.4%, *n* = 5 DSS-treated mice, 488 cells; 43.4 ± 3.3%, *n* = 4 formalin-injected mice, 859 cells) is shown. **(C)** The percentage of Brn3a-positive cells among c-fos cells (30.4 ± 2.6%, *n* = 5 DSS-treated mice, 1,141 cells; 18.1 ± 1.9%, *n* = 4 formalin-injected mice, 1,458 cells) and c-fos-H cells (38.1 ± 5.3%, *n* = 5 DSS-treated mice, 324 cells; 17.5 ± 1.6%, *n* = 4 formalin-injected mice, 807 cells) is shown. Horizontal bars indicate the median. Data were analyzed by the Mann-Whitney U test. ‘ns’ means non-significant. *p < 0.05.

### Characterization of Brn3a-Positive Neurons in the Spinal Dorsal Horn

Although the expression of Brn3a in the spinal dorsal horn has long been known (14), the identity of Brn3a-positive neurons is yet to be characterized. Thus, we assessed the expression of several molecular markers in Brn3a-positive neurons, including Pax2, preprodynorphin (Pdyn), and cholecystokinin (CCK) (Figure 5). Brn3a-positive neurons were rarely co-localized with Pax2 (Figure 5A-C, 0.7%, *n* = 2 mice, 158 cells) both in the marginal and the deeper laminae, suggesting they were excitatory neurons. Pdyn, which marks a population of excitatory and inhibitory neurons in lamina I and II (31), was positive in 45.4 ± 6.1% of Brn3a-positive neurons in the marginal lamina (Figure 5D-F, *n* = 3 mice, 251 cells). Furthermore, CCK, a marker of a population of excitatory neurons in lamina III and IV (32), was positive in 50.9% of Brn3a-positive neurons in deeper laminae (Figure 5G-I, *n* = 2 mice, 186 cells) but was not expressed in Brn3a-positive neurons localized in the marginal lamina (0%, *n* = 2 mice, 35 cells). These results suggest that Brn3a-positive neurons in the marginal lamina are excitatory neurons and neurochemically different from Brn3a-positive neurons localized in the deeper laminae.

**FIGURE 5.**
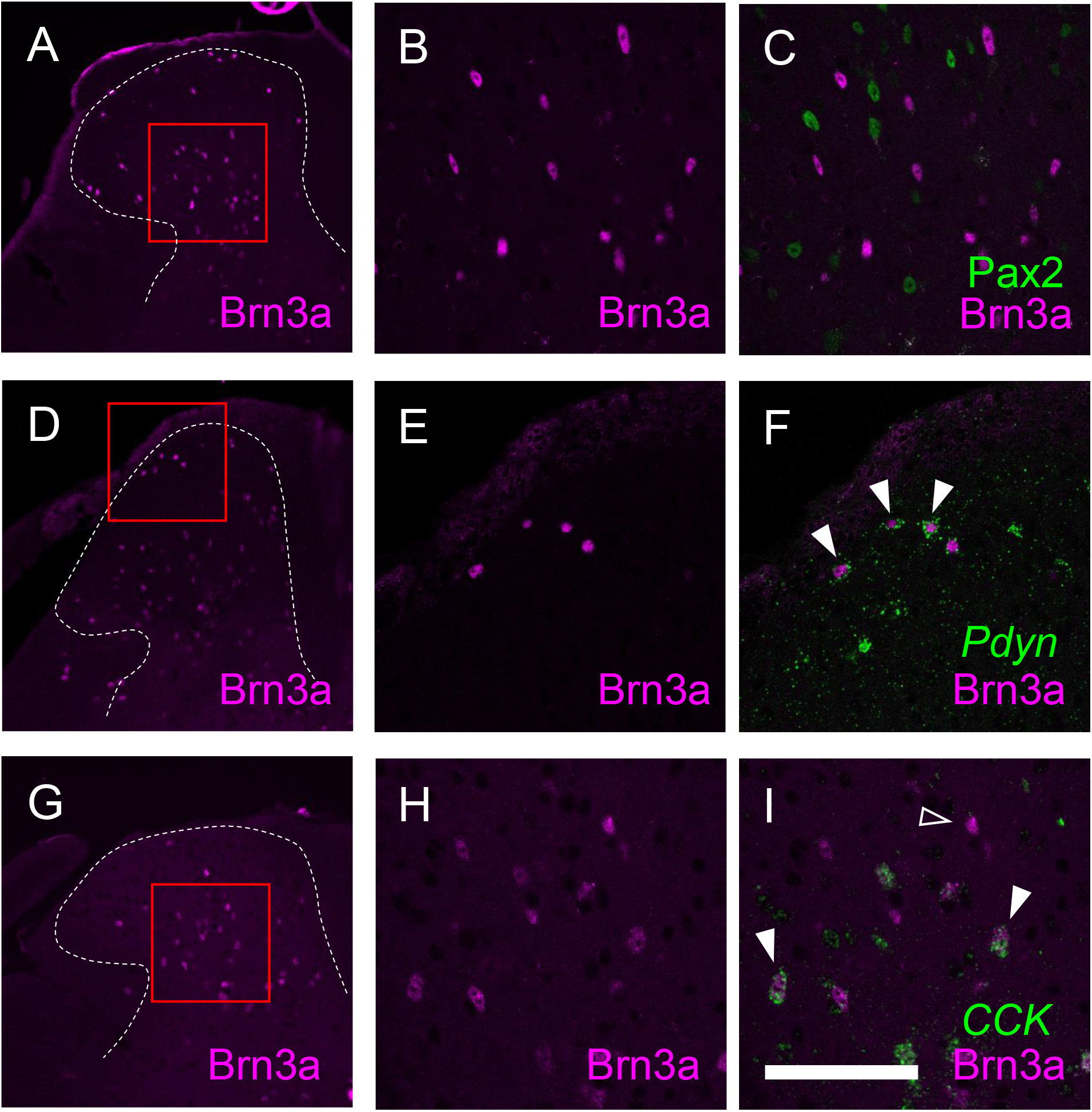
Characterization of Brn3a-positive neurons in the spinal dorsal horn by co-immunostaining with several markers. Brn3a immunostaining together with Pax2 immunostaining, *preprodynorphin* (*Pdyn*), or *cholecystokinin* (*CCK*) *in situ* hybridization was performed on transverse sections of the lumbosacral spinal dorsal horn of wild-type mice. **(A, D, G)** Low magnification images of Brn3a immunostaining (magenta) on the left spinal dorsal horn are shown. White dotted lines indicate the boundary between white and gray matters. **(B, C)** High magnification views (marked in **A**) of Brn3a signal with **(C)** or without **(B)** Pax2 signal (green) around the deep dorsal horn are shown. **(E, F)** High magnification views (marked in **D**) of Brn3a signal with **(F)** or without **(E)** *Pdyn* signal (green) around the shallow dorsal horn are shown. **(H, I)** High magnification views (marked in **G**) of Brn3a signal with **(C)** or without **(B)** *CCK* signal (green) around the deep dorsal horn are shown. Closed arrowheads indicate cells with positive for Brn3a and markers, whereas open arrowhead indicates Brn3a-single-positive cell. Scale, 100 μm.

Since supraspinal projection neurons that transmit nociceptive information are highly accumulated in the marginal lamina (33), we wondered whether Brn3a-positive neurons are localized in this area, including supraspinal projection neurons. We first focused on spino-parabrachial neurons innervating the lateral parabrachial nucleus (LPb) in the pons, which constitutes approximately 90% of total supraspinal projection neurons transmitting nociceptive signal`s` (33). These neurons were labeled by injection of the retrograde axonal tracer cholera toxin B (CTB) into the LPb (Supplementary Figure 5), and the spinal cord of injected mice was immunostained with antibodies against CTB together with Brn3a to analyze their co-localization (Figure 6A-C). Contrary to our expectations, the percentage of Brn3a-positive neurons among spino-parabrachial projection neurons in the marginal lamina was low at the lumbar level (14.5 ± 0.6%, *n* = 3 mice, 172 cells). We next examined spino-solitary projection neurons (34) (Supplementary Figure 5), since their axonal target, the NTS, is involved in the processing of visceral sensory information, although their visceral inputs have been assumed to be transmitted via the vagus nerve (35). Interestingly, the percentage of Brn3a-positive neurons among spino-solitary projection neurons in the marginal lamina was 51.8 ± 4.6% at the lumbar level (Figure 6D-G, *n* = 6 mice, 166 cells), which was significantly higher than that among spino-parabrachial projection neurons (*p* = 0.0238, Mann-Whitney U test). Similar results were obtained in the thoracic spinal cord of the mice, in which 16.8 ± 2.5 % (*n* = 3 mice, 97 cells) of spino-parabrachial projection neurons in the marginal lamina were Brn3a-positive versus 59.0 ± 9.9 % (*n* = 3 mice, 71 cells) of spino-solitary projection neurons being Brn3a-positive (Supplementary Figure 6).

**FIGURE 6.**
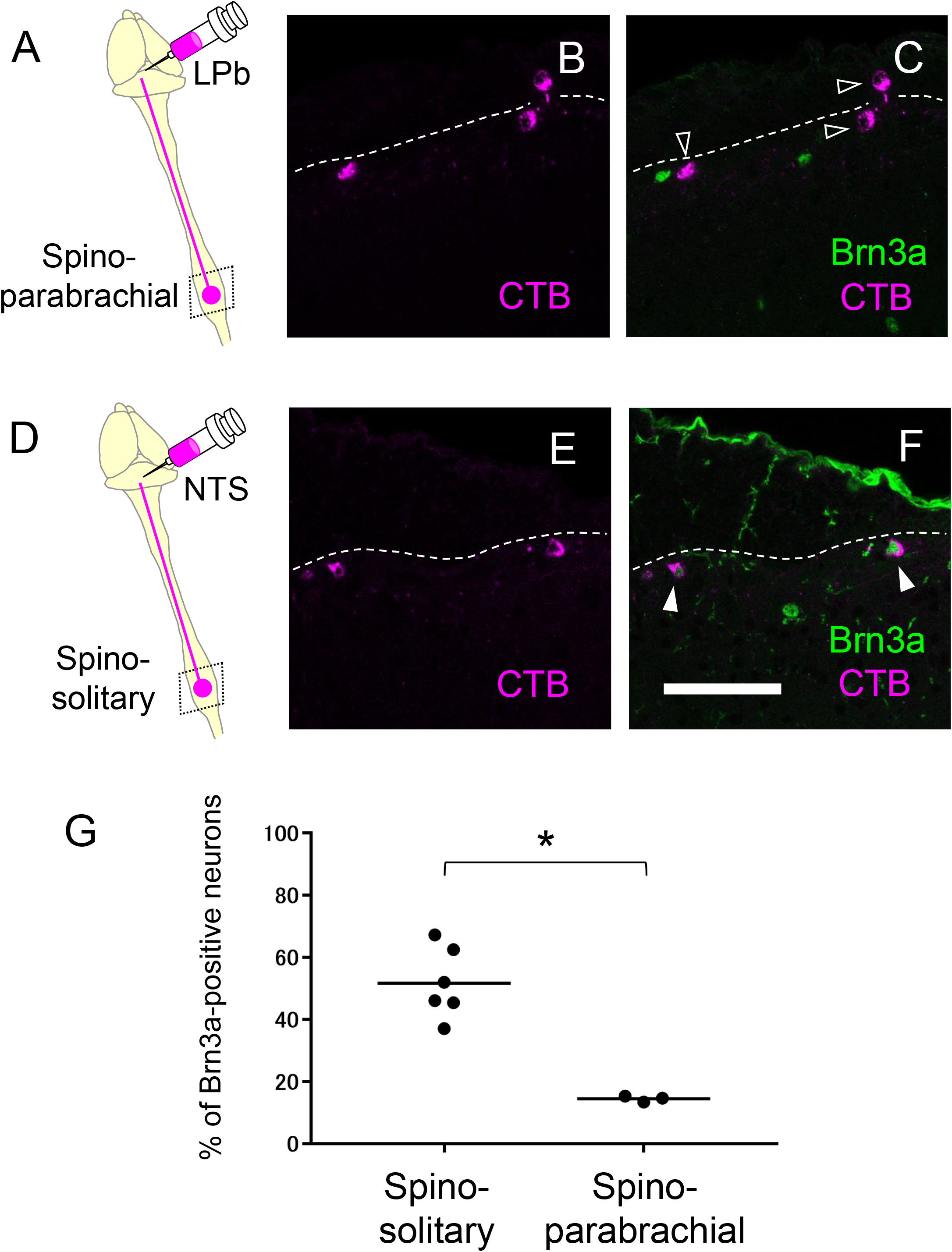
Expression of Brn3a in the spino-parabrachial and spino-solitary projection neurons. Spino-parabrachial **(A-C)** and spino-solitary **(D-F)** supraspinal projection neurons were labeled by injection of cholera toxin B (CTB) into the lateral parabrachial nucleus (LPb) and the nucleus of the solitary tract (NTS), respectively. Immunostaining of CTB (magenta, **B, C, E, F**) together with Brn3a (green, **C, F**) was performed on transverse sections of the lumbar spinal dorsal horn of the mice. White dotted lines indicate the boundary between white and gray matters in the shallow dorsal horn. Closed and open arrowheads indicate CTB- and Brn3a-double and CTB-single-positive cells, respectively. Scale, 100 μm. **(G)** Percentage of Brn3a-positive cells among spino-solitary (51.8 ± 4.6%, *n* = 6 mice, 166 cells) and spino-parabrachial (14.5 ± 0.6%, *n* = 3 mice, 172 cells) supraspinal projection neurons is shown. Horizontal bars indicate the median. Data were analyzed by the Mann-Whitney U test. *p < 0.05.

Preferential activation of Brn3a-positive spinal dorsal horn neurons in DSS-treated mice (Figure 3), together with enrichment of Brn3a in the spino-solitary projection neurons (Figure 6), implies that visceral pain transmission through these projection neurons strongly relies on the Brn3a-positive population. To examine this possibility, we labeled spino-solitary projection neurons with CTB in DSS-treated mice to assess the percentage of Brn3a-positive neurons among visceral pain-responsive spino-solitary projection neurons (Figure 7). In accordance with the abundance of Brn3a-positive neurons in the spino-solitary projection neurons (Figure 6), 57.4 ± 4.7 % (*n* = 3 mice, 57 cells) of c-fos- and CTB-double-positive neurons in the marginal lamina were Brn3a-positive. Collectively, these results suggest that a population of Brn3a-positive neurons in the marginal lamina of the spinal dorsal horn contributes to the transmission of visceral pain to the NTS.

**FIGURE 7.**
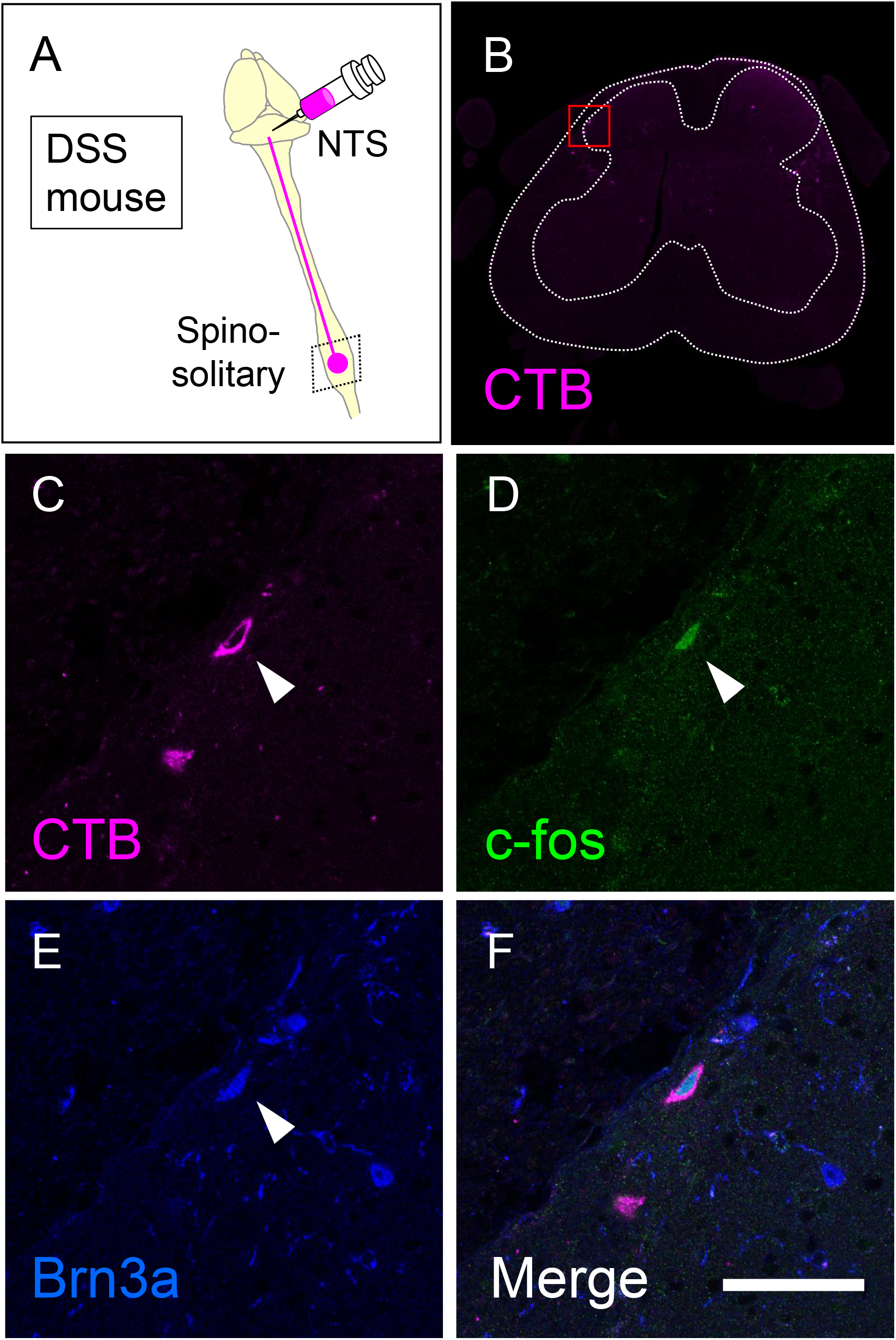
Expression of Brn3a in visceral pain-responsive spino-solitary projection neurons. **(A)** Spino-solitary supraspinal projection neurons were labeled by injection of cholera toxin B (CTB) into the nucleus of the solitary tract (NTS) in DSS-treated mice. Immunostaining of CTB together with c-fos and Brn3a was performed on transverse sections of the lumbosacral spinal cord of the mice. **(B)** A low magnification image of CTB immunostaining (magenta) on the representative section is shown. White dotted lines indicate the outline of the spinal cord and the boundary between white and gray matters. **(C-F)** High magnification views (marked in **B**) of CTB (**C** and **F**, magenta), c-fos (**D** and **F**, green), and Brn3a (**E** and **F**, blue) immunostaining are shown. Closed arrowheads indicate CTB-, c-fos- and Brn3a-triple-positive cells. Scale, 50 μm.

## DISCUSSION

In this study, we took advantage of IBD model mice generated by DSS treatment and analyzed visceral pain transmission in the spinal dorsal horn. Analyses of neural activity in the spinal dorsal horn of these mice demonstrated that visceral pain preferentially evokes the activity of Brn3a-positive neurons for transmission. We further revealed that a population of Brn3a-positive neurons in the marginal lamina was engaged in visceral pain transmission to the NTS.

### DSS-Treated Mice as a Visceral Pain Model

Neural circuits in the spinal dorsal horn that transmit visceral pain have been less studied compared to those of somatic sensation, possibly because of the lack of proper experimental models and difficulties in evaluating behavioral responses. Previously, DSS-treated mice have been employed in some studies on visceral pain in the colon, as DSS treatment enhanced visceral pain sensitivity to several stimulants injected into the colon (36, 37). Our study went a step further and showed that DSS treatment itself was sufficient to evoke neuronal activity in the SDH (Figure 2, Supplementary Figure 3) as well as visceral nociceptive behavior (Figure 1, Supplementary Figure 2).

The c-fos-positive neurons were found exclusively in the lumbosacral spinal cord (L6-S1) of DSS-treated mice (Figures 2 and 3 and Supplementary Figure 3). This result is in contrast to a previous study in which noxious mechanical stimulation of the colon triggered neural activity in both the lumbosacral and thoracolumbar (T10-L1) spinal cords (38). The difference was due to the stimulation of different layers of the colon in these models. The overall structure of the mucosa in the distal colon was severely damaged in DSS-treated mice without obvious anatomical changes in the other layers (Supplementary Figure 1). The sensation of the colorectal mucosa is mainly mediated through pelvic afferents, which innervate the lumbosacral spinal cords (27). This is consistent with the distribution of c-fos in the lumbosacral spinal cord of DSS-treated mice (Figures 2 and 3 and Supplementary Figure 3). In contrast, mechanical distention of the colon distorts not only the mucosa but also the surrounding layers such as the muscle and mesentery. The sensation of the muscle and mesentery is transferred through both splanchnic and pelvic afferents and eventually transmitted to not only lumbosacral but also thoracolumbar cords (27). DSS treatment induced c-fos expression in the lumbosacral cords, without additional mechanical stimulation of the colon (Figures 2 and 3 and Supplementary Figure 3). It is possible that inflammatory stimulants secreted in DSS-treated mice directly stimulated nociceptive afferents in the colon. Alternatively, innocuous mechanical stimulation by bowel movements may trigger noxious sensations in DSS-treated mice. Indeed, mechanically insensitive afferents, which account for 20% of all colorectal primary sensory afferents, were shown to become mechanically sensitive following the application of inflammatory stimulants (39).

DSS-treated mice exhibited an increased number and duration of anal licking of their anus (Figure 1). This behavior likely compensates for visceral pain caused by colitis for several reasons. First, the rectum, which is close to the anus, seems to be the most sensitive area among the colon in the perception of mechanical and chemical stimulation, because the expression of TRPV1, a critical mediator of visceral pain, is significantly higher in the rectum than in the distal, transverse, and proximal colon (40). Second, the central projection of primary sensory neurons responsible for the skin of the anus (∼S1) partially overlapped with the noxious sensory inputs in DSS-treated mice (L6-S1; Figure 2, Supplementary Figure 7).

### Involvement of Brn3a-Positive Spinal Dorsal Horn Neurons in the Transmission of Visceral Pain-Specific Signal

Our study demonstrated that the percentage of Brn3a-positive neurons among visceral pain-responsive neurons was significantly higher than that of somatic pain-responsive neurons (Figure 3U). This is in line with the preferential localization of visceral pain-responsive neurons in the marginal lamina of the SDH (Figure 2D-G). These results suggested that visceral pain preferentially activates Brn3a-positive neurons for transmission. Considering a previous study showing that almost all visceral pain-responsive neurons also receive somatic sensory inputs (4), it is highly possible that Brn3a-positive neurons responsive to visceral pain stimulation are also viscerosomatic neurons. This might explain why a small percentage of c-fos-positive neurons in formalin-injected mice were also Brn3a-positive (18.1±1.9%, Figure 3U). The absolute number of Brn3a- and c-fos-double-positive neurons in formalin-injected mice was comparable to that in DSS-treated mice (data not shown). Hence, it is conceivable that activation of viscero-somatic Brn3a-positive neurons would mainly contribute to the induction of visceral pain-related sensation, while activation of these neurons together with neurons responsive only to somatic pain (e.g., lamina II neurons) would induce somatic pain sensation. However, we cannot exclude the possibility that some populations of Brn3a-positive neurons respond only to visceral or somatic stimulation. Further electrophysiological or calcium imaging studies of Brn3a-positive neurons following visceral and somatic stimulation are needed to characterize the identity of these neurons.

### Visceral Pain Transmission to the Higher Brain Center

The current study demonstrated that Brn3a-positive neurons were highly represented in spino-solitary projection neurons responsive to visceral pain stimulation, although we did not characterize other populations of visceral pain-evoked Brn3a-positive neurons in the spinal dorsal horn (Figure 8). One possibility is that they might belong to propriospinal neurons that interconnect distant spinal segments. In support of this notion, most neurons expressing Brn3a and related transcription factors (Brn3b and Brn3c) in the central and peripheral nervous systems are long-range neurons (12). Thus, it is possible that Brn3a-positive neurons in the spinal dorsal horn are involved in long-range sensory transmission, including both the supraspinal and propriospinal pathways.

**FIGURE 8.**
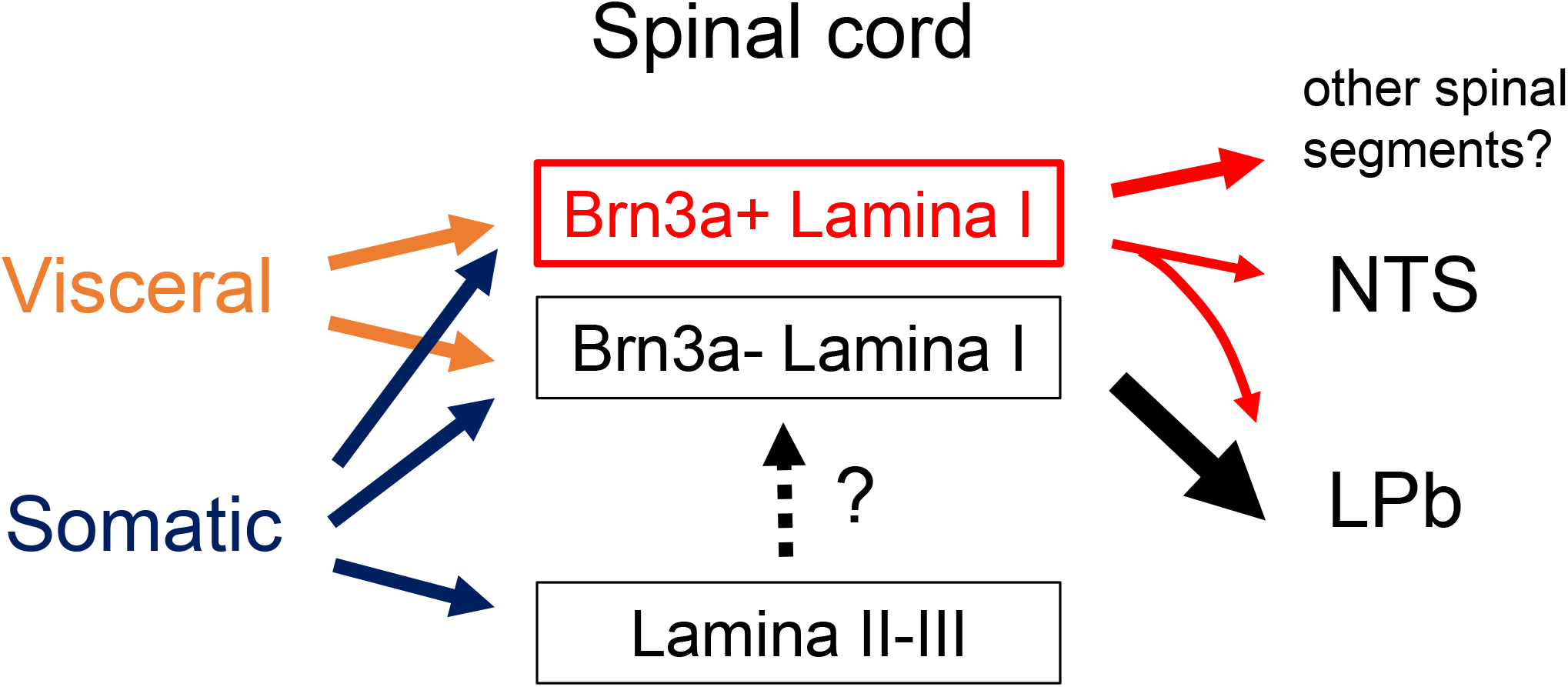
Schematic showing proposed neural circuits in the shallow dorsal horn for visceral and somatic pain. Visceral pain preferentially evokes neuronal activities of marginal (lamina I) lamina neurons in the spinal dorsal horn whereas somatic pain does those of both marginal and deeper laminae neurons (lamina II and III). Brn3a-positive neurons in the marginal lamina are one of the populations which preferentially responded to visceral pain. These neurons were highly represented among supraspinal projection neurons innervating NTS (spino-solitary projection neurons) compared to those innervating LPb (spino-parabrachial projection neurons). Note that most spino-solitary projection neurons also innervate LPb (41).

Among the supraspinal projection neurons examined in this study (Figure 6G), Brn3a-positive neurons were highly enriched in spino-solitary neurons (51.8 ± 4.6%) but not in spino-parabrachial projection ones (14.5 ± 0.6%), even though lateral parabrachial nucleus (LPb) is the principal axonal target of supraspinal projection neurons (33). Almost all (∼90%) spino-solitary projection neurons were shown to send collateral projections to LPb, whereas projection neurons innervating both NTS and LPb accounted for only a quarter of spino-parabrachial projection neurons (41)(Supplementary Figure 8). This fact suggests that supraspinal projection neurons can be largely divided into two populations in terms of projection toward the NTS and LPb: one innervates both the NTS and LPb and the other only the LPb. Based on the abundance of these neurons (41), the percentage of Brn3a-positive neurons among the latter population (spino-parabrachial only neurons) can be estimated to be 3.67% (Supplementary Figure 8), further indicating that Brn3a-positive neurons are enriched in projection neurons innervating both NTS and LPb compared to those innervating only LPb. Simultaneous activation of the NTS and LPb by Brn3a-positive projection neurons might be crucial for visceral pain transmission.

The NTS sends axonal projections to many brain areas, such as the ventrolateral medulla, parabrachial nucleus, periaqueductal gray, hypothalamus, and amygdala (42). Based on its connectivity, the NTS is commonly assumed to have an autonomic-affective function. Cardiovascular regulation of the NTS through the control of sympathetic premotor neurons in the rostral ventrolateral medulla has been well documented (43). Tachycardia evoked by nociceptive stimulation is shown to be suppressed by synaptic inhibition of NTS neurons (44), raising the possibility that visceral pain signaling via the spino-solitary pathway is involved in the regulation of cardiac function. On the other hand, NTS innervation to the amygdala suggests a role for the spino-solitary pathway in the transmission of an emotional aspect of visceral pain.

Since the vagal afferent nerve is an important mediator of viscerosensory transmission to the NTS (45), it may function as an integration center for visceral information through vagal and spino-solitary afferents. In addition, the involvement of PSDC neurons, which are localized outside the shallow dorsal horn, in the visceral pain transmission has long been known (10, 11). The PSDC neurons send axonal projections directly toward the gracile nucleus in the medulla, the lesion of which attenuates certain types of visceral pain. To fully understand visceral pain transmission, we need to elucidate the function of each of these pathways, as well as the mechanisms that integrate their information in the higher brain center.

## ACKNOWLEDGMENTS

We thank Drs. N. Funatsu, S. Iki, A. Inoue, T. Katano, K. Okuda, and R. Suno for their technical advice; Drs. N. Funatsu, A. Inoue, and T. Katano for materials; and Dr. Y. Zhu and T. Katano for critical reading of the manuscript. We would like to thank Editage (www.editage.com) for English language editing.

## FUNDING

This study was supported by a grant-in-aid (19K07855) for scientific research from the Ministry of Education, Science, Sports, and Culture of Japan.

## CONFLICT OF INTEREST STATEMENT

The authors declare no conflicts of interest associated with this manuscript.

## AUTHOR CONTRIBUTIONS

KN designed the study, performed the experiments, analyzed the data, and wrote the manuscript. SM contributed analytical tools and stereotactic surgery. TK contributed to the discussions and suggestions.

## Notes

### Competing Interest Statement

The authors have declared no competing interest.

### Summary of Updates

Figure 1 revised Figure 3 revised Supplemental files updated

