## Supplementary figure legends for "Involvement of Brn3a-positive spinal dorsal horn neurons in the transmission of visceral pain in inflammatory bowel disease model mice"

**Supplementary Figure 1. Histological analysis of the colorectal mucosa.**

(A) Hematoxylin and eosin (HE) staining was performed on the proximal (A, B), medial (C, D), and distal (E, F) colon of control (A, C, E) and DSS-treated (B, D, F) mice (10 µm in thickness). Scale, 100 µm.

**Supplementary Figure 2. Relationship between disease development and licking behavior of DSS-treated mice.**

(A) The severity of colitis was assessed based on body weight loss, diarrhea, and occult blood in the stool. The disease activity index (DAI) is the sum of the three parameters. (B, C) DAI scores of two DSS-treated mice (#1, and #2) from days 0 to 6 are shown. (D, E) The duration of the licking behavior of the mice during a 30 min time window is shown.

**Supplementary Figure 3. Global distribution of c-fos-positive cells in DSS-treated and control mice.**

The c-fos immunostaining of the transverse sections of the spinal cord from the upper thoracic to lower sacral segments in control (left) and two DSS-treated (center and right) mice was performed, and the distribution of c-fos-positive cells in 10 sections (20 µm in thickness, 3.2 mm apart) is shown. Cells with c-fos-intensity above the set threshold were considered c-fos-positive and are shown as red dots. The white and gray matter in each section are shown in light yellow and gray, respectively. Scale, 500 µm.

**Supplementary Figure 4. c-fos-positive cells in the lumbosacral spinal cord in formalin-injected mice.**

(A) Distribution of c-fos-positive cells. c-fos immunostaining on the transverse sections of the lumbosacral spinal cords in formalin-injected mice was performed, and distribution of c-fos-positive cells on six sections (20 µm in thickness, 800 µm apart) from the lumbar (~L5, top) to the sacral (~S2, bottom) spinal cords in a representative mouse in each condition is shown. The c-fos-positive cells are shown as red dots. White and gray matters in each section are shown in light yellow and grey colors, respectively. Formalin was injected into the lower back skin on the right (see materials and methods in detail). Scale, 500 µm. (B) The spinal cord was divided into the shallow dorsal horn (SDH), deep dorsal horn (DDH), ventral horn (VH), and lateral funiculus (LF). The distribution of c-fos-positive cells in these areas in formalin-injected mice (*n* = 4 mice, 942 cells) is shown in the pie chart. (C-F) Transverse sections of the spinal dorsal horn of formalin-injected mice were immunostained with anti-c-fos (magenta) together with anti-PKCγ (green; E, G) antibodies. Images on the right dorsal horn around L6 (D, E) and S1 (F, G) segments are shown. Scale, 100 µm.

**Supplementary Figure 5. CTB injection into the NTS and LPb.**

Immunostaining using an anti-CTB antibody was performed on coronal sections of the hindbrain in mice in which CTB was injected into the NTS or LPb on the right. (A-D) The CTB immunostaining of hindbrain sections of mice injected with CTB into the NTS. The NTS region (circled by dashed red lines) was identified using dark field photographs. (E-H) CTB immunostaining of hindbrain sections of mice injected with CTB into the LPb. The LPb lies dorsal and lateral to the superior cerebellar peduncle (SCP dashed red line), which was identified using dark field photographs. The outline of the brain is indicated by the dashed white line. Anterior to the left. Dorsal up. Scale, 1 mm. CTB, cholera toxin B; NTS, nucleus of the solitary tract; LPb, lateral parabrachial nucleus.

**Supplementary Figure 6. Percentage of Brn3a-positive neurons among supraspinal projection neurons at the thoracic level.**

Spino-solitary and spino-parabrachial supraspinal projection neurons were labeled by injection of cholera toxin B (CTB) into the nucleus of the solitary tract (NTS) and the lateral parabrachial nucleus (LPb), respectively. Immunostaining of CTB and Brn3a was performed on transverse sections of the thoracic spinal dorsal horn of the mice. Percentage of Brn3a-positive cells among spino-solitary (59.0 ± 9.9%, *n* = 3 mice, 71 cells) and spino-parabrachial (16.8 ± 2.5%, *n* = 3 mice, 97 cells) supraspinal projection neurons is shown. Horizontal bars indicate medians.

**Supplementary Figure 7. Distribution of c-fos-positive cells and the central projection pattern of primary sensory neurons responsible for the skin of the anus.**

Primary sensory neurons responsible for the transmission of innocuous mechanical stimulation of the anal skin on the left side were labeled with cholera toxin B (CTB). The mice were subsequently treated with water containing 2% DSS for seven days. The distribution of c-fos (green) and CTB (magenta) in the transverse sections of the lumbosacral spinal cord (L6 to S2) of mice was analyzed by immunostaining. The ipsilateral side was on the left side. Scale, 500 µm.

**Supplementary Figure 8. Percentage of Brn3a-positive neurons among spino-parabrachial projection neurons which do not send collaterals to the nucleus of the solitary tract.**

Estimation was based on a previous study on double labeling of spino-solitary and spino-parabrachial neurons [33], together with the current study (Fig. 5G).
