## Supplementary figures for "Involvement of Brn3a-positive spinal dorsal horn neurons in the transmission of visceral pain in inflammatory bowel disease model mice"

### Sup Figure 1

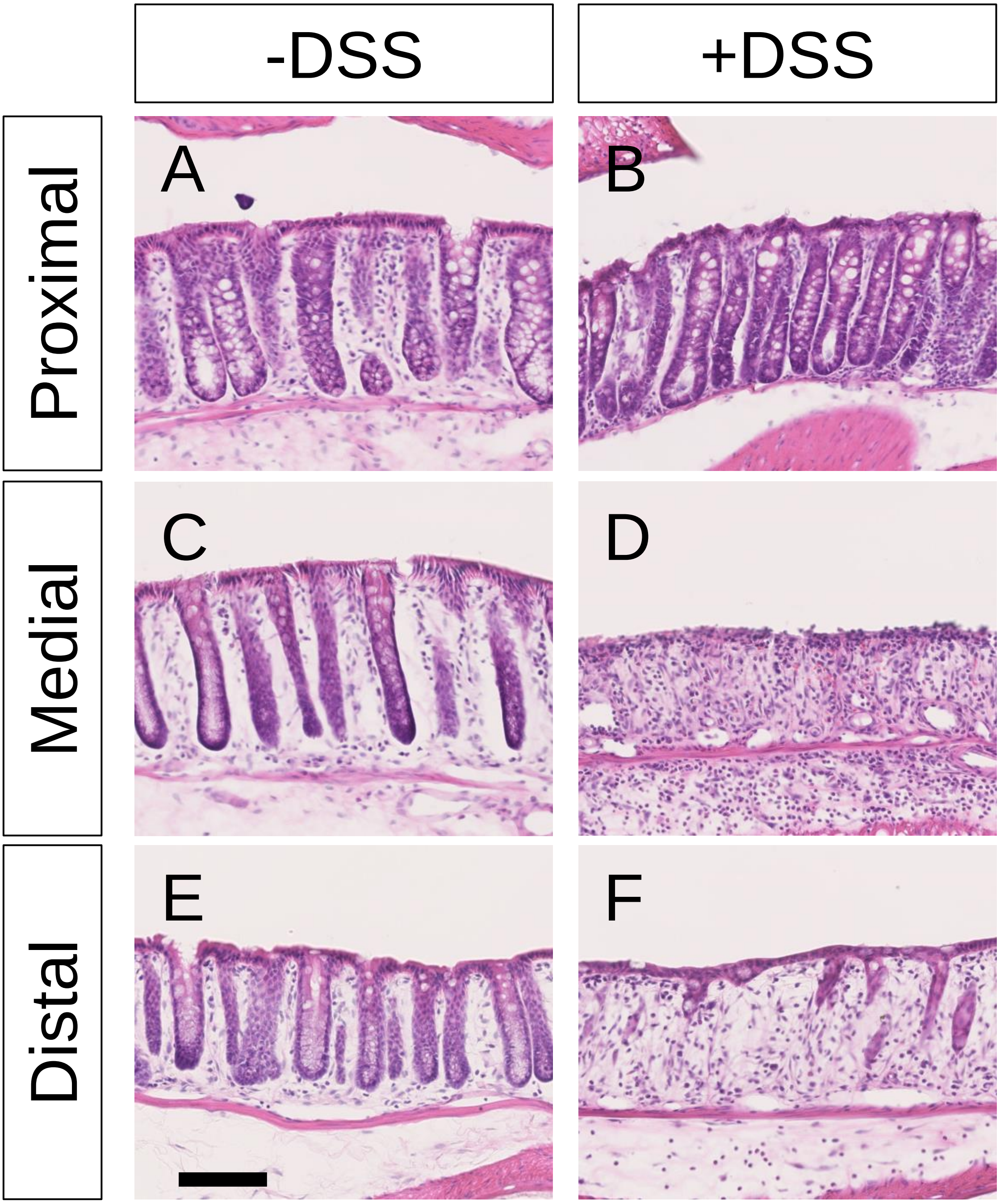

### Sup Figure 2

#### A: DAI score

| Score | Weight loss | Stool | Blood |
|---|---|---|---|
| 0 | None | Normal | None |
| 1 | 0-5% | Adhere to animal bedding | |
| 2 | 5-10% | Shape became round | Slightly bleeding |
| 3 | 10-15% | Shape collapse | |
| 4 | 15-20% | Watery | Gross bleeding |

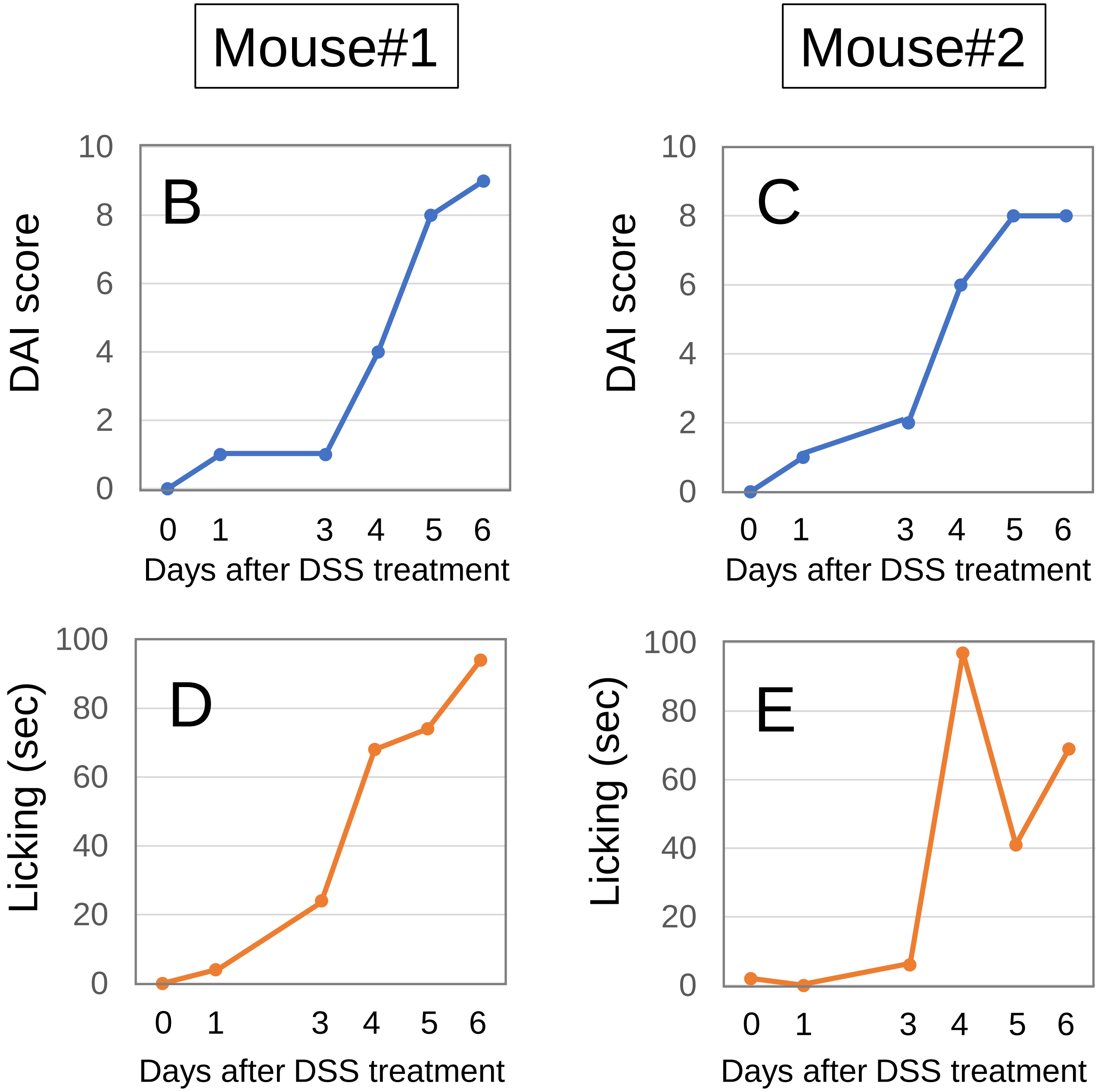

### Sup Figure 3

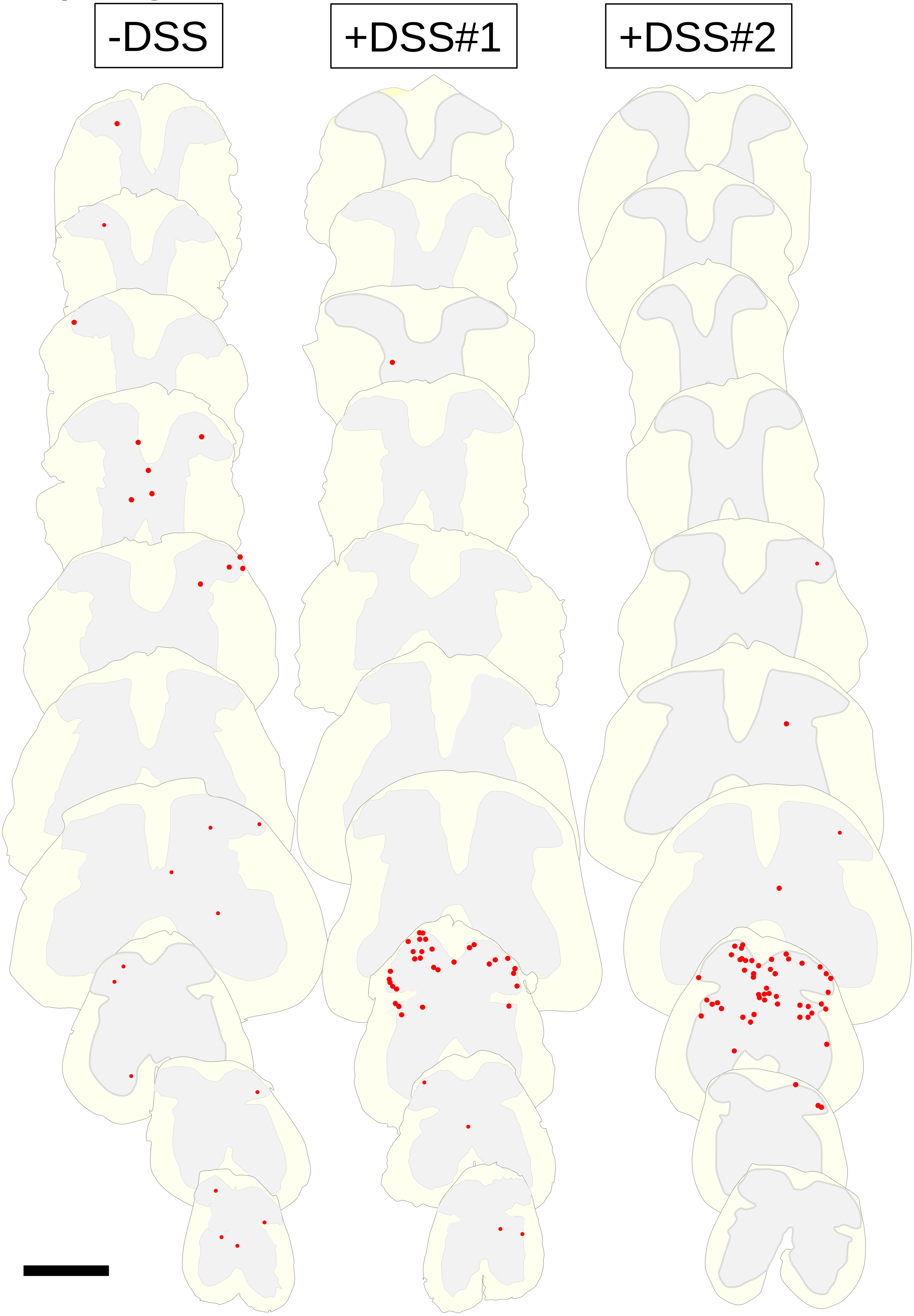

### Sup Figure 4

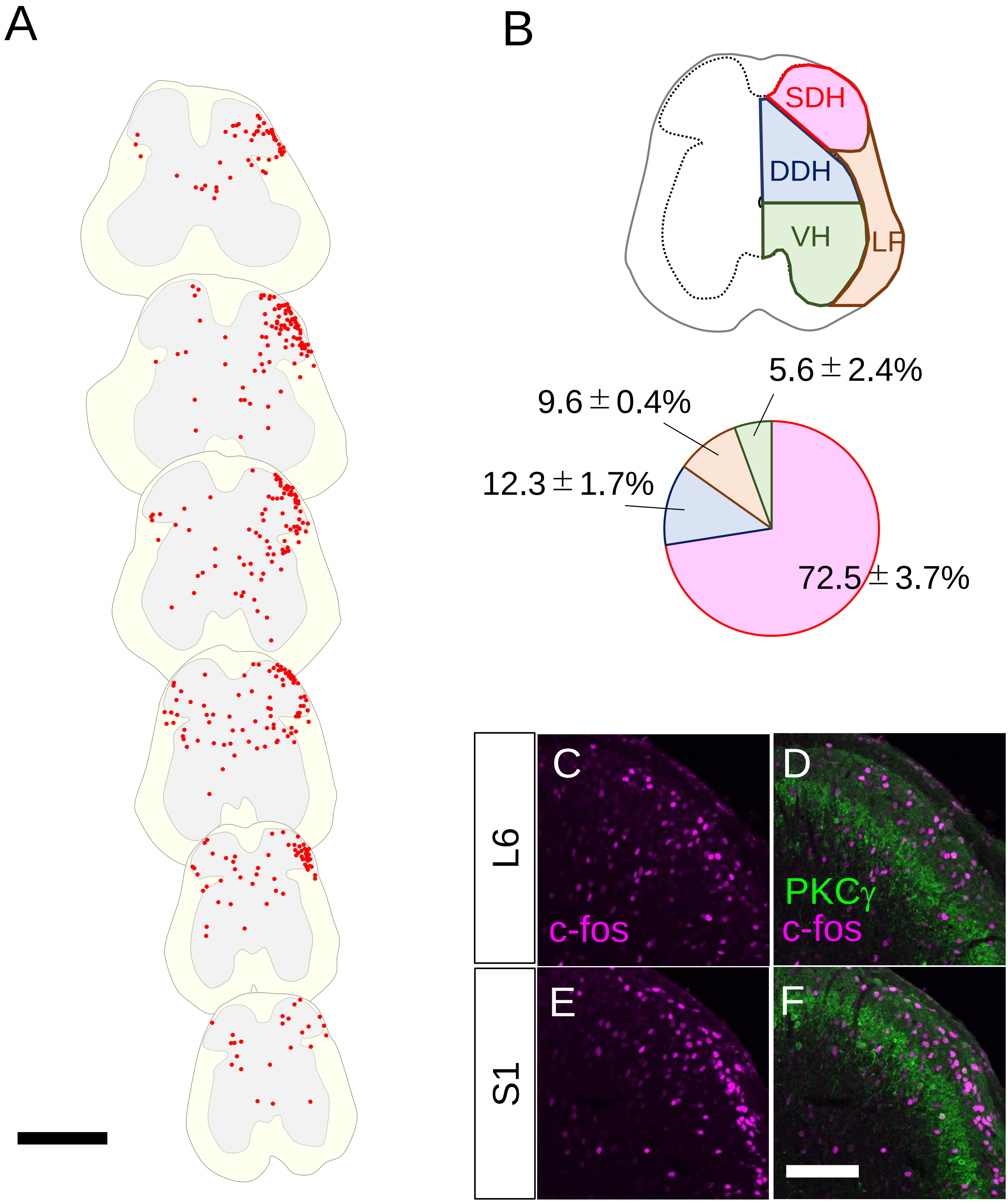
A
B
SDH
DDH
VH
LF
5.6±2.4%
9.6±0.4%
12.3±1.7%
72.5±3.7%
L6
S1
C
D
E
F
c-fos
PKCγ
c-fos

### Sup Figure 5

CTB → NTS

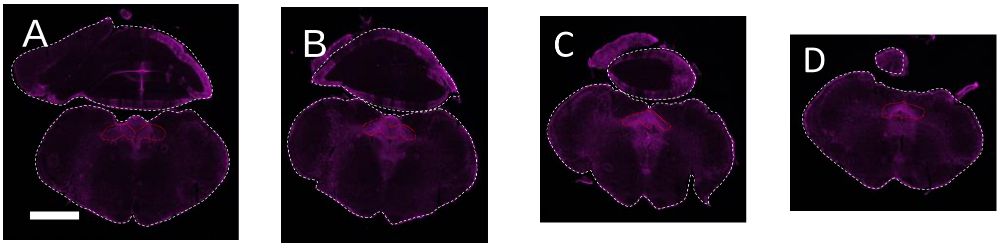


240 μm

CTB → LPb

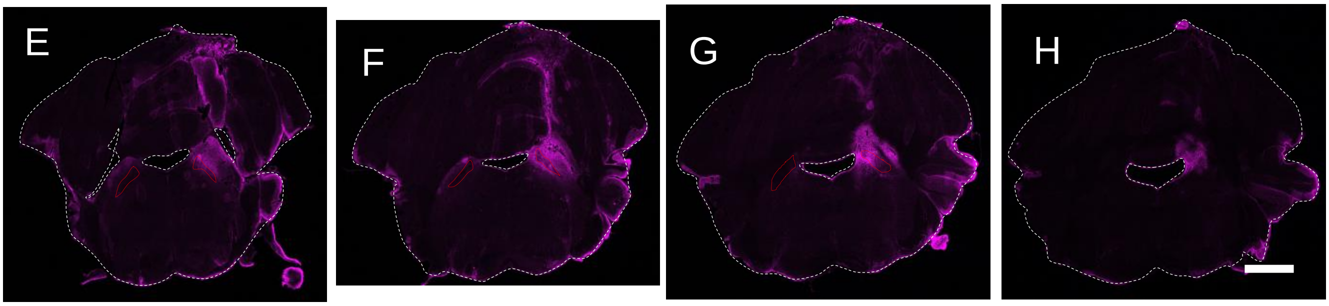


160 μm

### Sup Figure 6

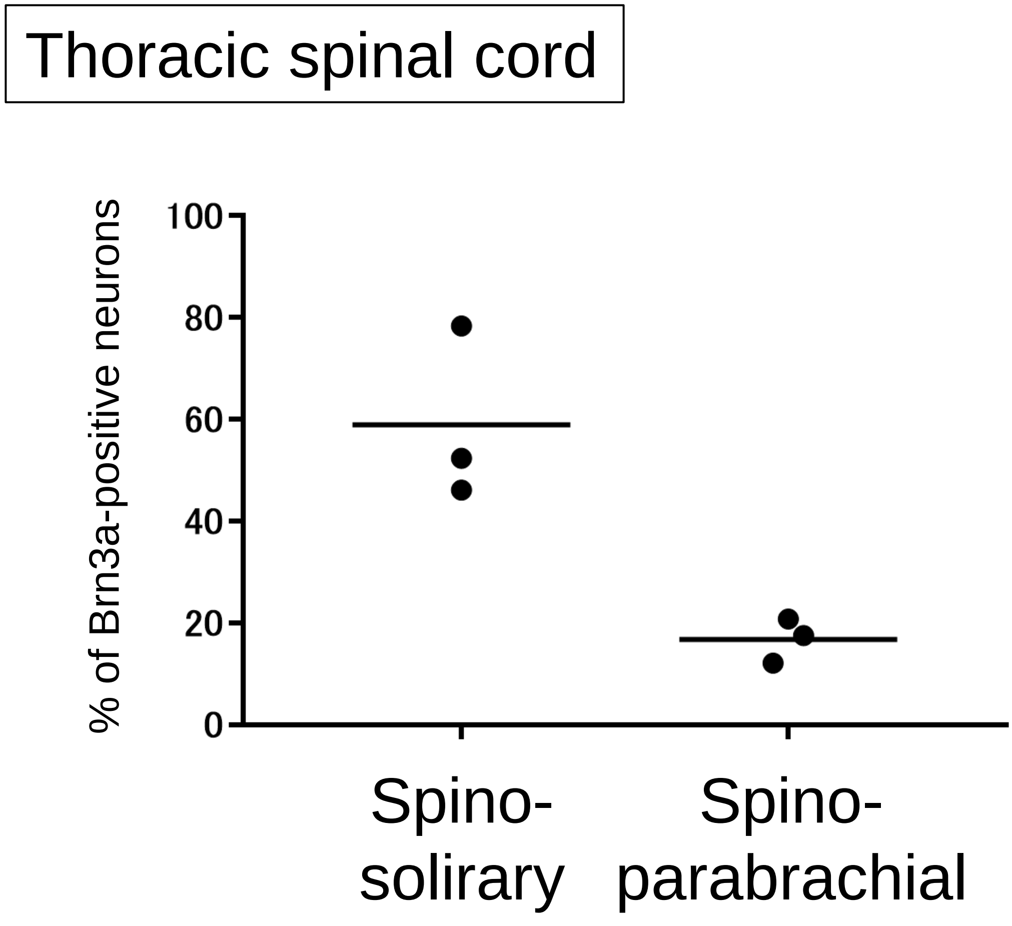
Thoracic spinal cord
% of Brn3a-positive neurons
100
80
60
40
20
0
Spino-
solirary
Spino-
parabrachial

### Sup Figure 7

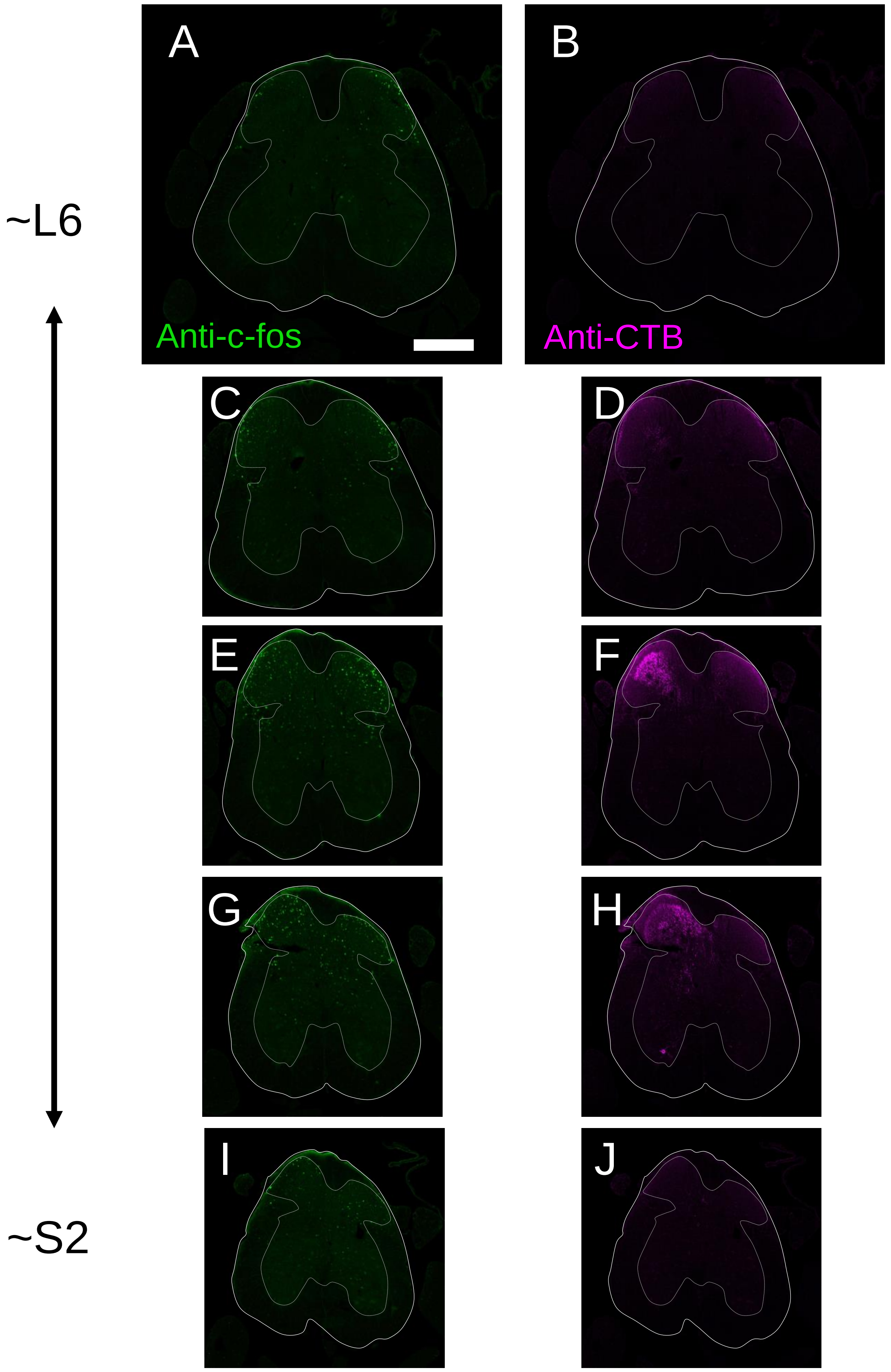

### Sup Figure 8

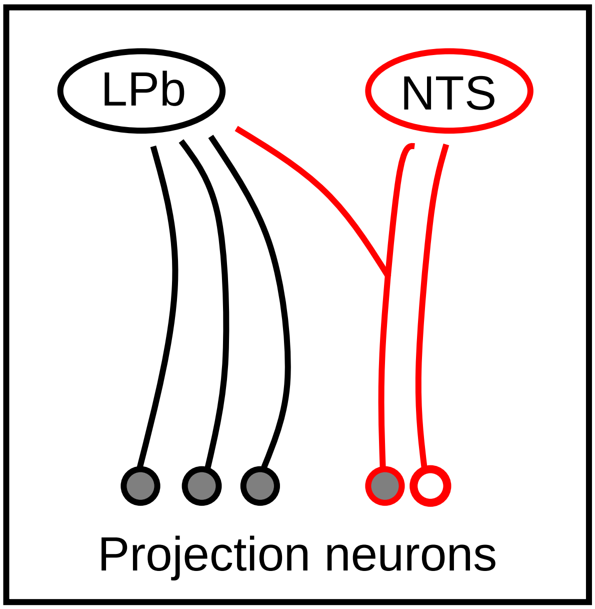


Polgar et al (2010) showed that ……

ratio of LPb projection neurons ( + ) to NTS projection neurons ( + ) is 4:1

ratio of LPb+NTS projection neurons ( ) to NTS only projection neurons ( ) is 9:1

The current study showed that ……

percentage of Brn3a positive neurons among LPb projection neurons ( + ) is 14.5%

percentage of Brn3a positive neurons among NTS projection neurons ( + ) is 51.8%

If the number of NTS projection neurons ( + ) is N

the number of LPb projection neurons ( + ) is 4N,
and the number of Brn3a-positive LPb projection neurons is 4N x 0.145

The number of LPb+NTS projection neurons ( ) is 0.9N,
and the number of Brn3a-positive LPb+NTS projection neurons is 0.9N x 0.518

The number of LPb only projection neurons ( ) is 4N – 0.9N = 3.1N,
and the number of Brn3a-positive LPb only projection neurons is 4N x 0.145 – 0.9N x 0.518

Taken together, the percentage of Brn3a-positive neurons among LPb only projection neurons is

$$\frac{4N \times 0.145 - 0.9N \times 0.518}{3.1N} \times 100 = 3.67\%$$
